# PDBe CCDUtils: an RDKit-based toolkit for handling and analysing small molecules in the Protein Data Bank

**DOI:** 10.1101/2023.08.04.552003

**Authors:** Ibrahim Roshan Kunnakkattu, Preeti Choudhary, Lukas Pravda, Nurul Nadzirin, Oliver S. Smart, Qi Yuan, Stephen Anyango, Sreenath Nair, Mihaly Varadi, Sameer Velankar

## Abstract

While the Protein Data Bank (PDB) contains a wealth of structural information on ligands bound to macromolecules, their analysis can be challenging due to the large amount and diversity of data. Here, we present PDBe CCDUtils, a versatile toolkit for processing and analysing small molecules from the PDB in PDBx/mmCIF format.

PDBe CCDUtils provides streamlined access to all the metadata for small molecules in the PDB and offers a set of convenient methods to compute various properties using RDKit, such as 2D depictions, 3D conformers, physicochemical properties, scaffolds, common fragments and cross-references to small molecule databases using UniChem. The toolkit also provides methods for identifying all the covalently attached chemical components in a macromolecular structure and calculating similarity among small molecules. By providing a broad range of functionality, PDBe CCDUtils caters to the needs of researchers in cheminformatics, structural biology, bioinformatics and computational chemistry.

## Introduction

The Protein Data Bank (PDB) [1], managed by the worldwide PDB (wwPDB) consortium [2], serves as the single global repository for information on 3D structures of proteins, nucleic acids, and complex assemblies. With over 200,000 entries as of July 2023, about 75% of these structures contain at least one small molecule bound to a protein or nucleic acid. Some small molecules are present due to experimental necessities, such as aiding crystallisation [3–5] or enabling cryoprotection [6–8], while others play biologically significant roles acting as cofactors, metabolites, or drugs [9–12]. Information on ligand conformations and their interaction with macromolecular partners is crucial for deciphering their role in biological function and mechanisms.

To maintain standardised and accurate data on these small molecules, the wwPDB maintains Chemical Component Dictionary (CCD). This comprehensive reference resource contains data for all unique chemical components, including individual protein and nucleic acid residues and small molecules found in PDB entries [13]. CCD provides chemical description, composition, connectivity, and idealised coordinates for every unique chemical component.

During the wwPDB annotation process, each deposited structure undergoes careful processing to identify individual chemical entities, which are then compared against the already existing CCD to ensure consistent representation [14]. Despite playing a significant role as a reference dictionary, CCD description has certain limitations due to the nature of deposited data and wwPDB annotation practices. The wwPDB annotation guidelines suggest the CCD definition represents a neutral form for every compound [15]. In some cases, ligands may be refined and deposited as a set of covalently linked components, where each component represents a distinct CCD component. In other cases, ligands with peptide linkages may be split into the respective CCD components during annotation [14]. As a result of these structure determination practices or wwPDB annotation policies, ligands are fragmented into smaller components in almost 6% of PDB entries, limiting a straightforward interpretation of the complete ligand and its function. Due to such fragmentation of large ligands, some CCDs may not represent a chemically reasonable ligand on their own, leading to inaccuracies in representing the actual chemical structures of small molecules. These limitations can lead to incomplete or incorrect interpretation of ligand interactions and hinder mechanistic insights into protein function. To mitigate these issues, the wwPDB developed the Peptide-like molecules Reference Dictionary (PRD) specifically addressing the fragmentation issue for peptide-like inhibitors and antibiotic ligands [16]. Nevertheless, any other complex ligand in the PDB continues to be fragmented, preventing easy interpretation of the complete chemical structures and their interactions with macromolecules. Additionally, both CCD and PRD contain minimal data, lacking molecular properties and cross-references to relevant small molecule databases, making comprehensive analyses or integration of relevant information an arduous task. Another persistent challenge is a useful and accurate 2D depiction of CCDs to easily visualise the chemical structures and discern ligand characteristics.

Furthermore, the advances in structure determination techniques are resulting in an increasing number of structures for large macromolecular machines [17, 18]. The increasing number of depositions is also accelerating the number of new small molecules in the PDB archive and very shortly the traditional three-letter code will be exhausted, leading to a transition to five-letter CCD identifiers [19]. These developments will result in PDBx/mmCIF becoming the only source of information as both large structures or those that have five-letter CCD identifiers can not be represented in the legacy PDB format. To help smooth the transition to PDBx/mmCIF and the various challenges highlighted earlier, we have developed a versatile and user-friendly Python package, the PDBe CCDUtils, for handling and analysing small molecules in the PDB.

The core functionality of PDBe CCDUtils is based on RDKit, an open-source cheminformatics toolkit widely used in the scientific community for handling molecular data [20]. RDKit provides many features, including molecular structure manipulation, molecular descriptor calculation, and various cheminformatics algorithms. Building upon RDKit’s capabilities, PDBe CCDUtils extends its functionality specifically in the context of PDB data, allowing researchers to overcome the limitations faced with CCD and PRD. By utilising PDBe CCDUtils, researchers in cheminformatics, structural biology, bioinformatics, and computational chemistry can obtain an accurate representation of the small molecule and efficiently analyse the data in the PDB.

## Implementation

PDBe CCDUtils offers a comprehensive suite of functionalities that enhance the handling and analysis of small molecules in the PDB. By serving as a wrapper around essential RDKit functionality, PDBe CCDUtils seamlessly integrates the capabilities of RDKit with PDBx/mmCIF files, empowering users to access a wide array of cheminformatics tools and molecular manipulation techniques for small molecules in the PDB with ease and efficiency. The toolkit facilitates the reading and writing of small molecule reference PDBx/mmCIF files, automatically instantiating RDKit objects representing small molecules and their attributes, streamlining the process of accessing and analysing PDB data for further research.

PDBe CCDUtils also includes vital functionality to validate the chemical sanity of molecules, ensuring accurate representation and adherence to chemical rules, thereby preventing inaccuracies in ligand interactions analysis. The challenge of the 2D depiction of molecules is addressed by the toolkit, offering the option to generate the best 2D depiction using templates or connectivity. This ensures visually meaningful representations of chemical structures, enabling researchers to comprehend ligand characteristics accurately.

To overcome the fragmentation issue caused by multicomponent ligands being represented as separate CCD components in the PDB entries, PDBe CCDUtils introduces Covalently Linked Components (CLCs). By defining CLC molecules that encompass the entire set of individual CCD components which are covalently bonded together, PDBe CCDUtils provides a precise and chemically complete representation of these multicomponent ligands present in the PDB. PDBe CCDUtils also includes functionalities to find scaffolds and search common fragments/sub-structures against a fragment library, enabling researchers to explore structural similarities and analyse essential pharmacophoric elements in small molecules within the PDB. The implementation details of all these core functionalities are discussed in detail below.

### Parsing small molecules reference files to access meta-data and RDKit computed properties

PDBe CCDUtils supports input files in PDBx/mmCIF format, the master format for the PDB archive [21]. This data dictionary format is the basis of wwPDB data deposition, annotation, and archiving of PDB data from all supported experimental methods [21]. The wwPDB FTP area provides access to PDB’s small molecule reference files for CCDs and PRDs in PDBx/mmCIF format [22, 23]. Additionally, the PDBx/mmCIF format files for CLCs are provided in PDBe FTP area [24]. PDBe CCDUtils uses Gemmi, an open-source and efficient PDBx/mmCIF parser [25], for reading and writing these files. After reading the definitions of chemical components from small molecule reference files, they are represented as a Component object in the PDBe CCDUtils library, which is the core structural representation of a chemical component. The Component object is a wrapper around the “`rdkit.Chem.rdchem.Mol`’’ object. The Component object provides easy access to all metadata information encoded in the small molecule reference PDBx/mmCIF file or enriched later through the PDBeChem pipeline, an integral part of PDBe’s weekly release process [26], providing up-to-date and comprehensive small molecule information in the PDB (Figure 1). The CCDs/PRDs reference files include properties such as id, name, formula, pdbx_release_status, pdbx_modified_data, and descriptors including systematic chemical names and chemical descriptors (SMILES, InChI, InChIKey). The computed properties are generated using the structural data accessible from the `‘mol’` attribute of the Component object. The `‘mol’` attribute is a “`rdkit.Chem.rdchem.Mol`” object generated from the parsed atom coordinates and their connectivity in the reference file. CCD/PRD components also have ideal coordinates generated by Molecular Network’s Corina [27] or OpenEye’s OMEGA [28] and model coordinates from the PDB entry where the CCD/PRD component is first observed. Correspondingly, an Ideal conformer and Model conformer of type `“rdkit.Chem.rdchem.Conformer”` are added to the “`rdkit.Chem.rdchem.Mol`” object to access these coordinates.

**Figure 1.**
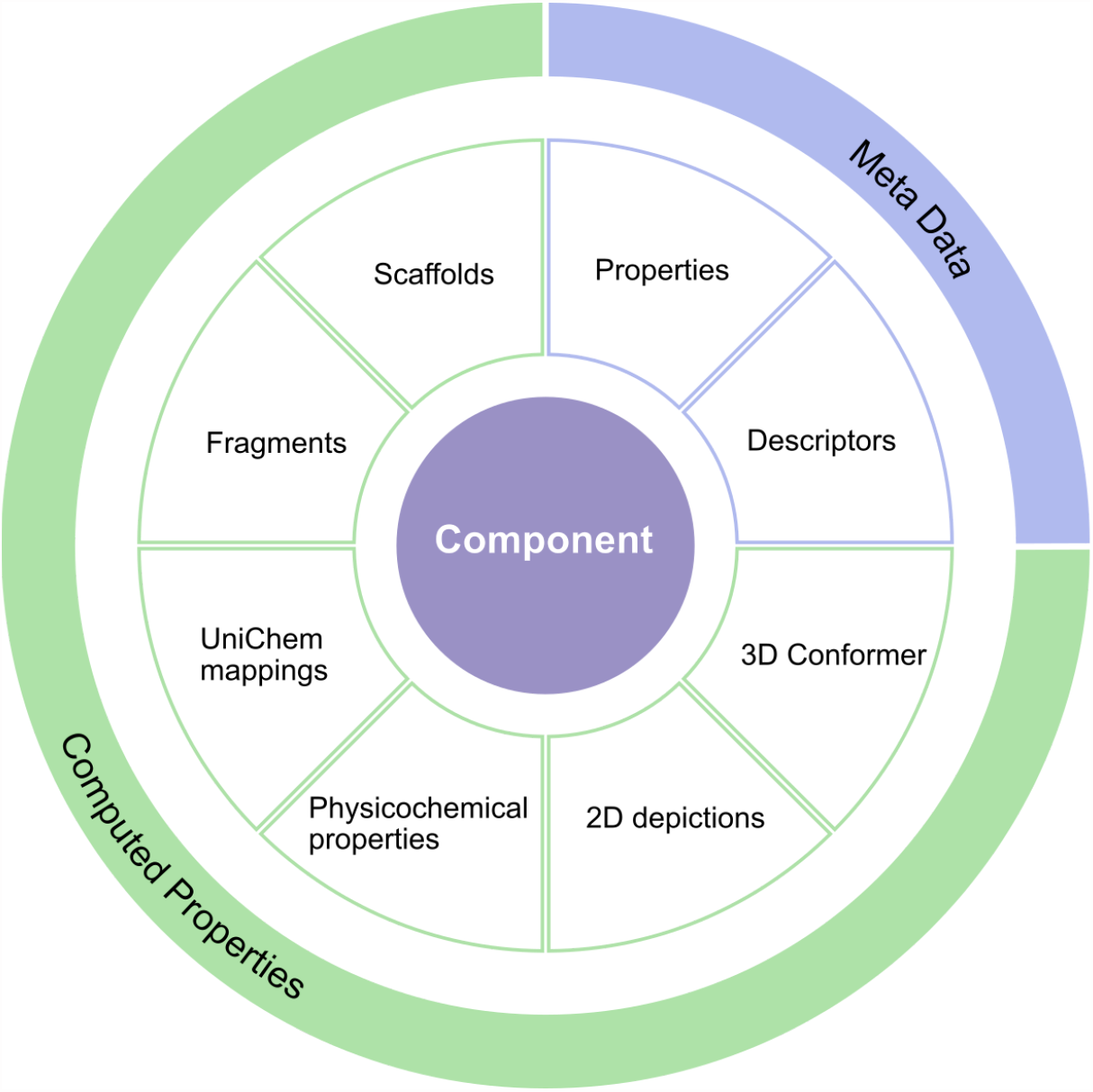
PDBe CCDUtils Component representation. The Component object is the core structural representation of small molecules in PDBe CCDUtils. It provides access to metadata such as descriptions and properties, and computed properties, such as scaffold and fragment information, cross-references from UniChem and physicochemical properties of small molecules.

After adding atom coordinates and their connectivity to the “`rdkit.Chem.rdchem.Mol object`”, an augmented sanitisation is carried out using RDKit as described below. Once a Component object is generated for a CCD/PRD, it can be exported to multiple formats. PDBe CCDUtils currently supports exporting to SDF, CIF, PDB, JSON, XYZ, XML and CML formats.

### Augmented data sanitisation process

PDBe CCDUtils also has an augmented data sanitisation process to address representation issues found in certain CCD files, ensuring efficient handling of biological ligands in PDB.

For instance, heme (CCD identifier HEM) contains a metalloporphyrin ring with two nitrogens atoms, each with a valency of 4. However, the default`‘rdkit.Chem.Sanitizemol’`function in RDKit fails to generate the complete representation of such ligands, with bonds to metal atoms, leading to unusual valency problems and incomplete structures. To overcome this limitation, PDBe CCDUtils implements an iterative sanitisation procedure. Initially, the molecule undergoes sanitisation using the “`rdkit.Chem.SanitizeMol`“ function, and any unusual valencies issues and affected atoms are identified. The augmented procedure then adjusts the bond type between the affected atom and a metal atom to be single while modifying the formal charges accordingly. This iterative procedure is repeated until there are no unusual valency issues reported by RDKit or a maximum of ten times. By resolving these representation issues, PDBe CCDUtils ensures the generation of accurate and chemically sensible molecules.

### Identifying covalently linked components

Several large ligands in the PDB are split into individual CCD components. For instance, PDB entry `6lq4` is a structure of Acyl-CoA dehydrogenase bound to its substrate Myristoyl-CoA. However, the substrate Myristoyl-CoA is split into Myristoyl (CCD ID: MYR) and CoA (CCD ID: COA). Such splitting of ligands into individual CCD components makes it difficult to correctly identify their biological relevance and their interactions with macromolecules and mapping to other small molecule databases. The Peptide-like molecules Reference Dictionary (PRD), tackles this fragmentation issue for a subset of peptide-like inhibitors and antibiotic ligands. We have addressed this issue for all the remaining cases by utilising PDBe CCDUtils and defined Covalently Linked Components (CLC) for ligands consisting of multiple covalently linked CCD components, providing a more precise and comprehensive representation of these multicomponent ligands in the PDB (Figure 2).

**Figure 2.**
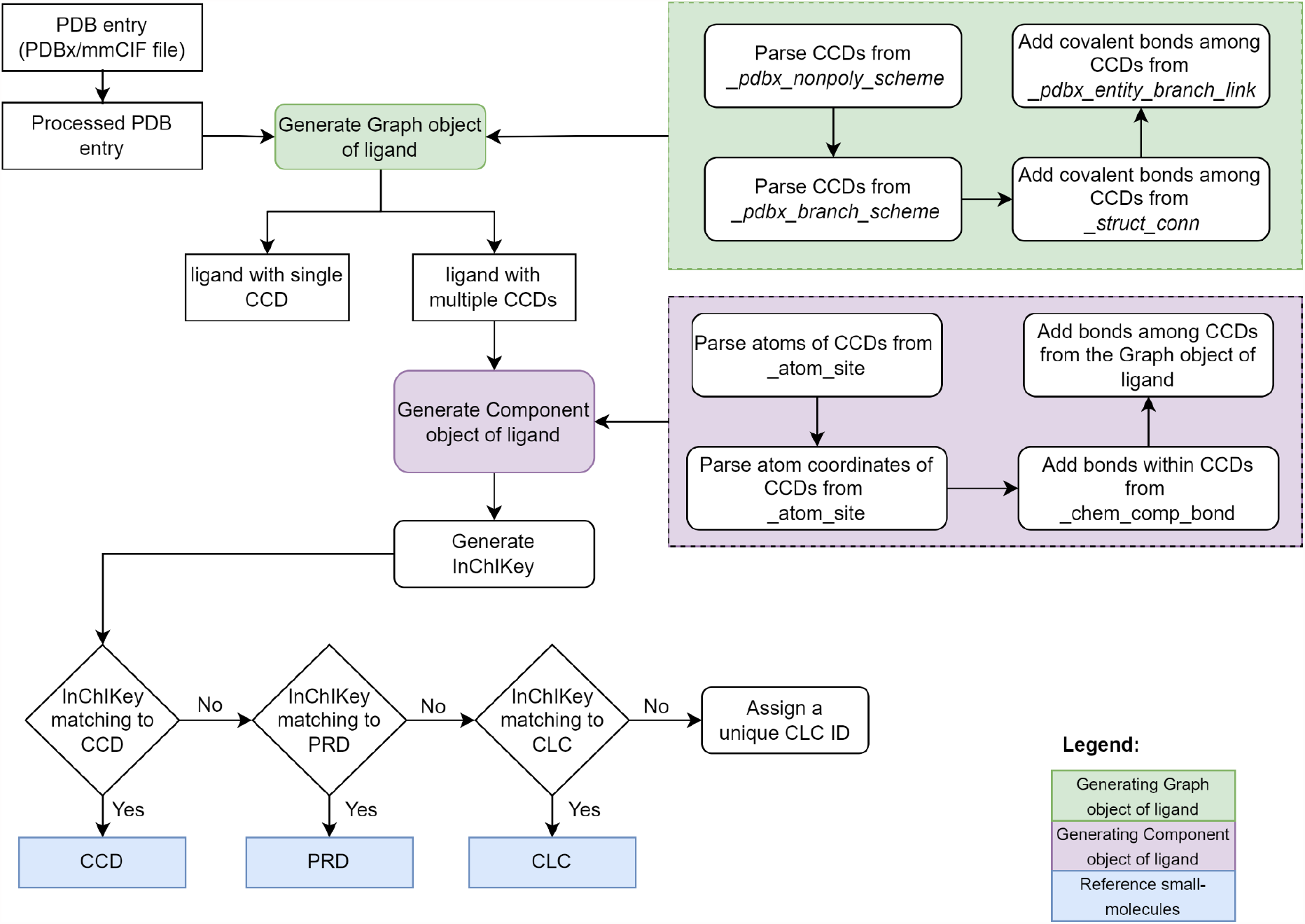
Schematic workflow of identifying unique CLCs. The method identifies Covalently Linked Components by processing a PDB structure file, generating a graph object of ligand with identified CCD components as nodes and covalent bonds as edges, and then converting the graph object into a Component object by parsing atom information and 3D coordinates, and adding bonds within CCD components from the processed PDB entry file and adding bonds among CCD components from the graph object of ligand.

Identifying CLCs starts by reading molecular structure data from an individual PDB entry file in PDBx/mmCIF format. The file is then processed to keep only the first model if there are multiple models, then only the atoms with maximum occupancy in case of alternate conformations are considered. Next, a graph object of ligand [29] is generated using CCDs identified from the “`_pdbx_nonpoly_scheme`” and the “`_pdbx_branch_scheme`” data categories as nodes and adding covalent bonds parsed from the “`_struct_conn`” and “`_pdbx_entity_branch_link`” categories as edges. If the ligand is composed of multiple CCDs linked by covalent bonds, then the graph object of the ligand is converted to a Component object in PDBe CCDUtils by parsing information about atoms and 3D coordinates from “`_atom_site`”, adding bonds within each CCD from “`_chem_comp_bond`” and adding bonds among CCDs from edges between nodes of the graph object described above. If the component’s InChIKey does not match any existing reference small molecules (CCDs, PRDs, or previously identified CLCs), it is classified as a new Covalently Linked Component (CLC). As part of the weekly PDBeChem pipeline at PDBe [26], this new CLC is assigned a unique identifier, facilitating identification and access to these molecules.

### 2D image generation of ligands in PDB

RDKit offers various methods for generating 2D depictions of ligands [30], but no single method is universally optimal for all the ligands in the PDB. Hence, we defined a score to quantify the quality of 2D depictions and select the best image generated by the template-based or connectivity-based methods of PDBe CCDUtils. The template-based method generates 2D depictions using a template molecule via the RDKit function “`rdkit.Chem.AllChem.generateDepictionMatching2DStructure”`. The template can be user-provided or downloaded from PubChem [31]. A hand-curated set of ten templates are also provided with PDBe CCDUtils which is available at [32]. If a user chooses to use a template from PubChem, PDBe CCDUtils uses the PubChem API [33] to download the template molecule based on the InChIKey match. The PubChem templates are rescaled to a bond length of 1.5 Å before generating depictions, as it is the default bond length of depictions generated by RDKit. The connectivity-based method uses the “`rdkit.Chem.rdCoordGen`” module to generate 2D coordinates.

PDBe CCDUtils runs both methods for all ligands, and a Depiction Penalty Score is calculated to determine the best image. We observed that the quality of the 2D depiction deteriorates when bonds collide and atoms crowd together in the 2D space. To account for this, we defined the Depiction Penalty Score (DPS) as a weighted sum of the number of bonds colliding and the number of pairs of atoms in suboptimal positions:

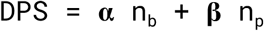

Where **α** is the bond collision penalty, n_b_ is the number of bonds colliding, **β** is the suboptimal atom position penalty, and n_p_ is the number of pairs of atoms closer to each other than the RDKit’s standard distance. We set **α** to 1 and **β** to 0.4 by default. A DPS of zero indicates a high-quality 2D depiction, while higher scores indicate lower quality. To count the number of bonds colliding, we considered two cases. First, when an atom is shared between two bonds, we consider the bonds to be colliding if the angle between them is less than 10°. Second, when no atoms are shared between two bonds, Cramer’s rule is used to check for collisions [34]. Similarly, we classified pairs of atoms to be in suboptimal positions if the distance between them is less than 0.5 Å. To consider such pairs of atoms in suboptimal positions, we counted the number of atoms within 0.5 Å radius of each atom using the “`spatial.KDTree.query_ball_point”` function from SciPy [35].

### Scaffolds and a curated library of fragments

Scaffolds are core chemical substructures characterising a group of molecules [36]. Their relevance is reflected in those compounds sharing the same scaffold are likely to have similar synthetic pathways [36]. Consequently, scaffolds with preferable biological activities are often used as initial templates for compound synthesis and diversification in small molecule drug discovery [37]. Although there are various definitions for molecular scaffolds, PDBe CCDUtils uses the most commonly used methods implemented in RDKit - MurckoScaffolds [38] and BRICS [39]. These algorithms can be easily accessed to generate scaffolds of small molecules in PDB via the `‘get_scaffolds’` method of the Component object. Fragments are small chemical structures that may contribute to binding to a target macromolecule [40]. In fragment-based drug discovery, the potency of small fragments with relatively weak binding affinities can be improved by chemically combining them to form larger structures with higher specificity and improved binding characteristics [40]. PDB ligands can be searched against a library of 2158 fragments manually curated by PDBe [26], ENAMINE [41] and Diamond-SGC-iNext Poised Library (DSiP) [42] using the `‘library_search’` method. Alternatively, an external library of fragments can also be supplied as a tab-delimited file in the same format as the default fragment library in PDBe CCDUtils, which is available at [43].

## Results and Discussion

The PDBe CCDUtils package provides a range of functionalities for analysing and manipulating small molecules in PDB structures. Here we provide examples of how PDBe CCDUtils can be used in a scientific context.

### Identification of CLCs in PDB entries

Figure 3 showcases four examples of Covalently Linked Components (CLCs) identified using PDBe CCDUtils in different PDB entries. These representations highlight the complete chemical structure of CLCs, which were previously fragmented into multiple CCD components during PDB deposition. Leveraging the InChIKeys of these CLCs, we were able to map and cross-reference these molecules to various external databases such as DrugBank, ChEMBL, ChEBI, and PubChem as shown in Table 1 and is available to the user in CIF files available at PDBe FTP area [24]. Such cross-references to other external databases facilitate easier access to valuable information about the biological and chemical contexts in which these molecules are found, providing researchers with deeper insights into their functional and pharmacological relevance.

**Table 1.**
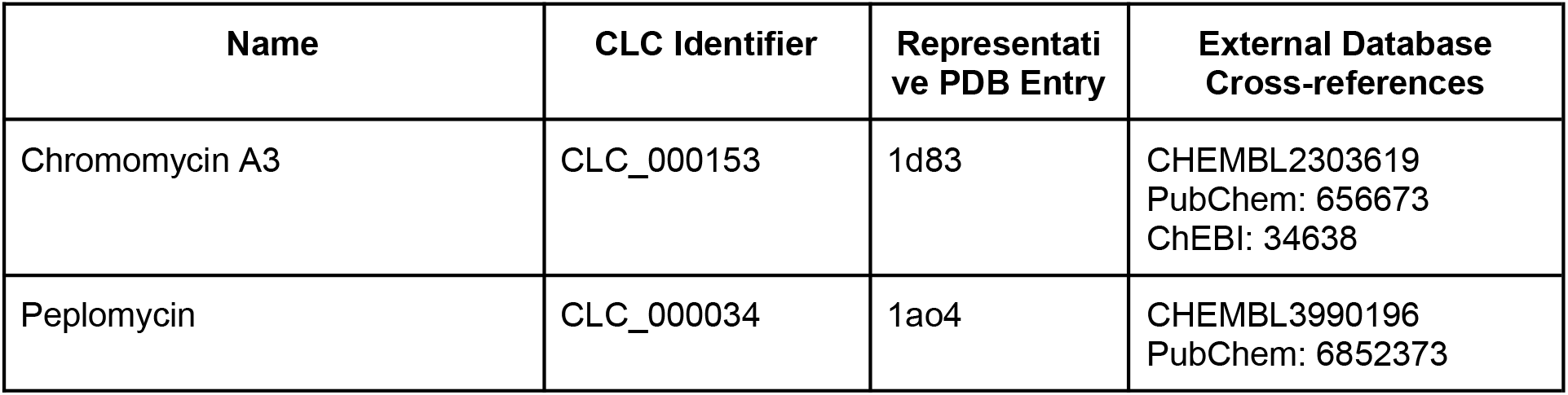
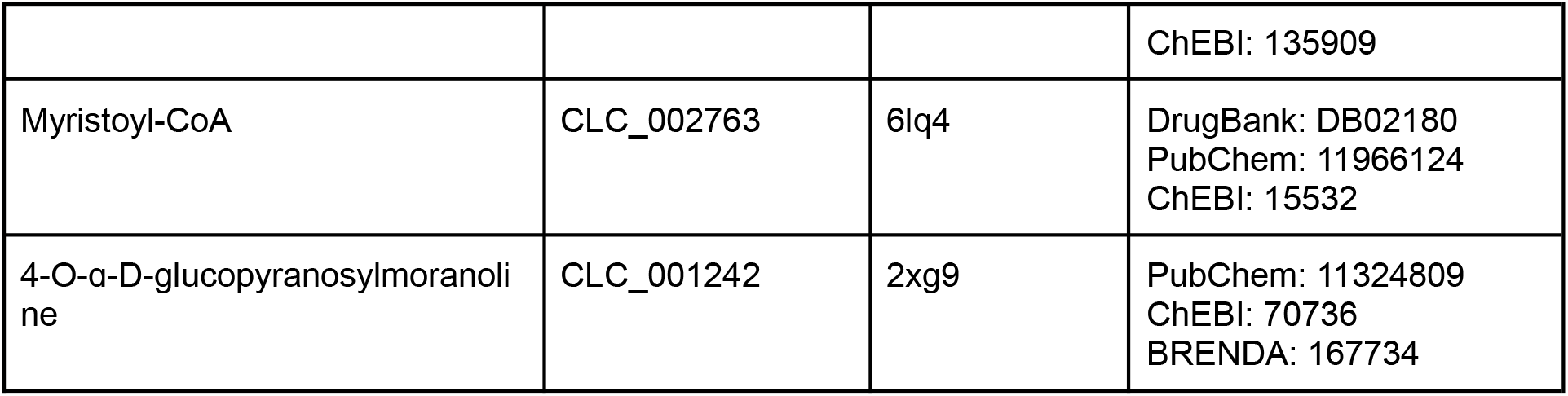
Examples of CLCs with cross-references to other external resources.

**Figure 3.**
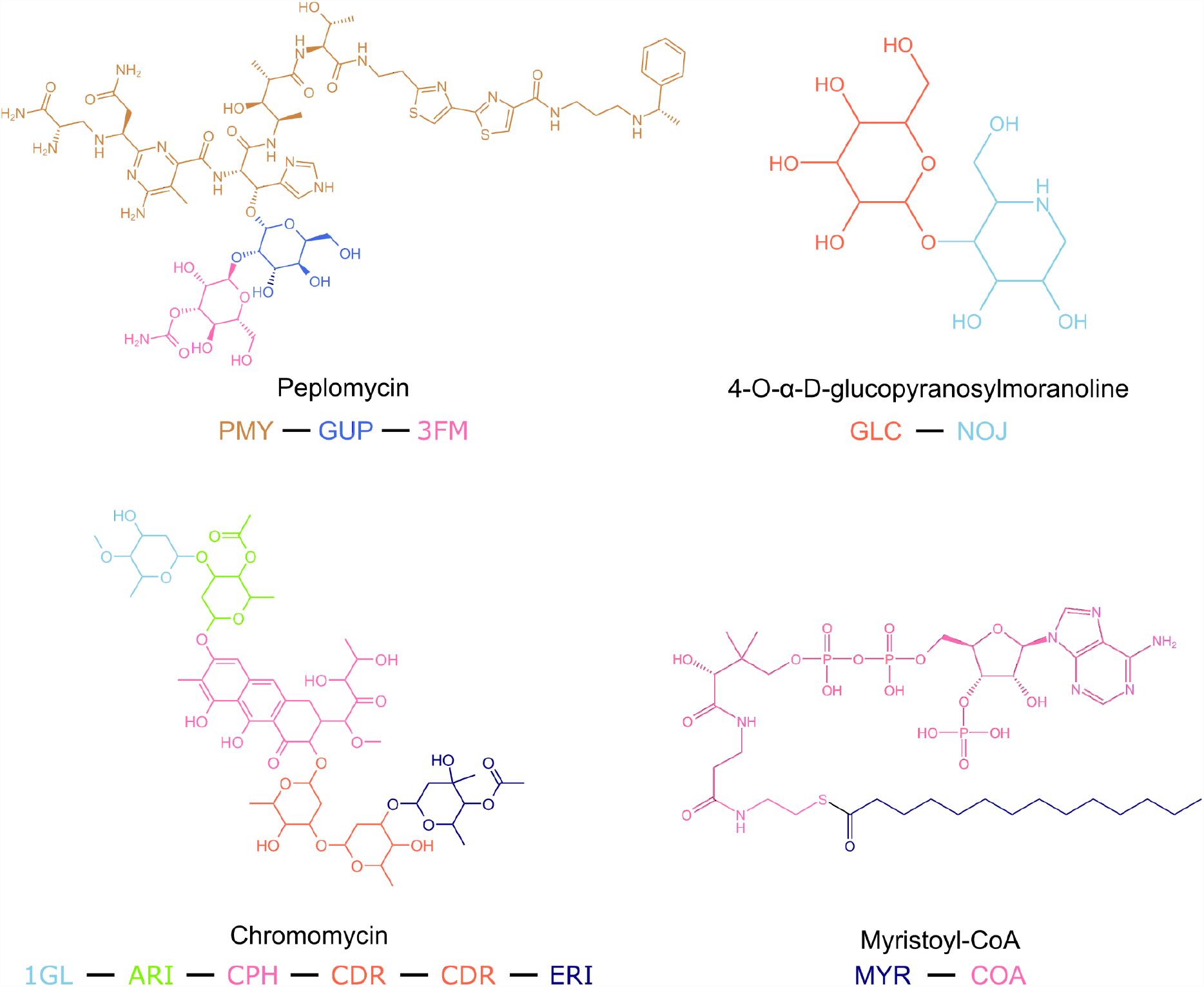
Examples of CLCs. Examples of Covalently Linked Components (CLCs), coloured based on their constituting chemical components (CCDs).

### Indication of quality of 2D images using the DPS

The DPS is a valuable metric to assess the quality of 2D depictions generated using RDKit [44]. A DPS of zero indicates that there are no bond clashes, and there is adequate distance between atoms in the 2D depiction, resulting in high-quality 2D representation. As the value of DPS increases, the quality of the 2D depiction deteriorates. Figure 4 presents an example of 2D depictions generated for the CCD component `‘HME’ (`porphycene containing iron) using the connectivity-based and the template-based methods in PDBe CCDUtils. The template-based method generates a high-quality 2D depiction without bond-clashes using the Porphycene template in PDBe CCDUtils. The best 2D image and 2D coordinates for each CCD, PRD and CLC based on the DPS is available from the PDBe FTP area [24].

**Figure 4.**
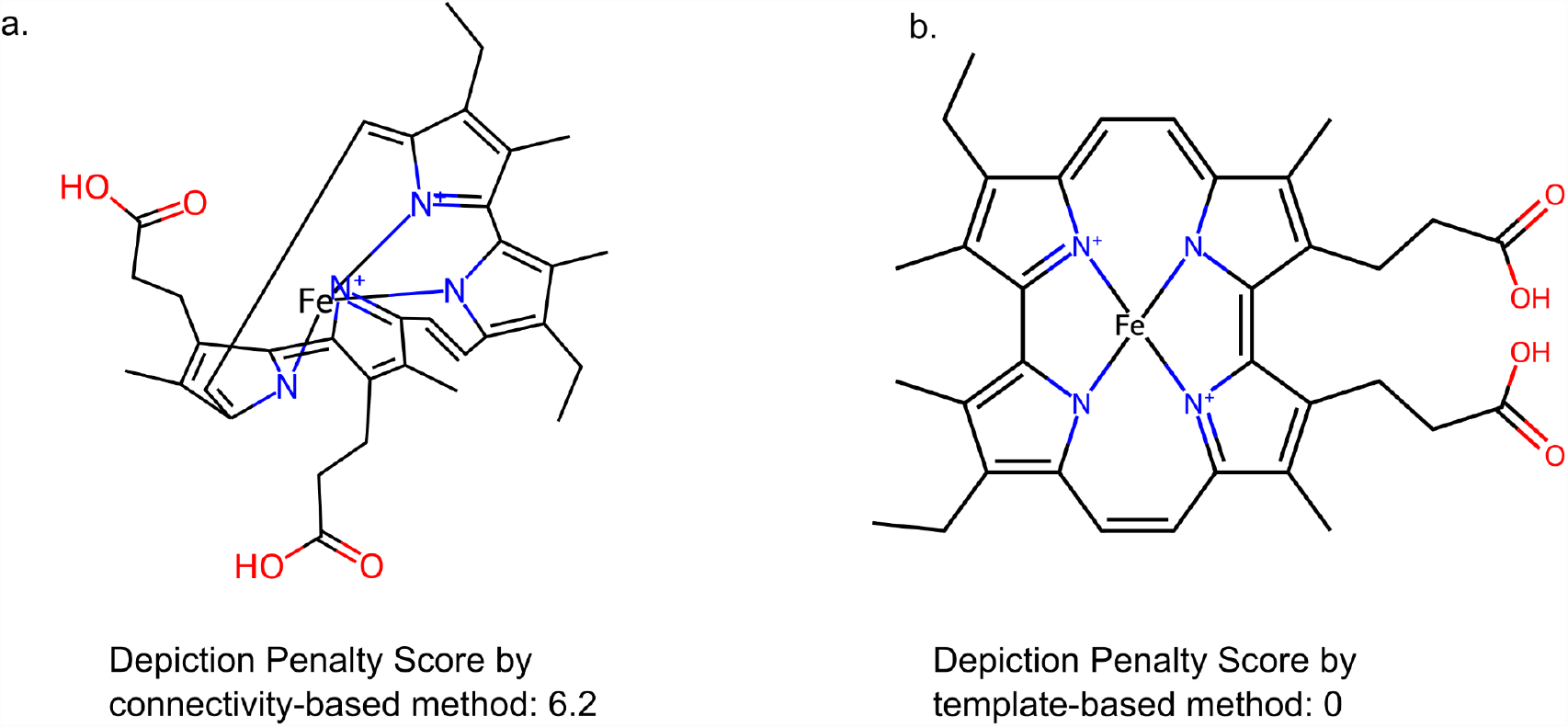
Example for use of Depiction Penalty Score. The Depiction Penalty Score is a metric to assess the quality of 2D depictions generated using RDKit. (a) 2D depiction of `‘HME’` with a high Depiction Penalty Score, highlighting some minor bond clashes and suboptimal atom distances in the 2D depiction, generated using the connectivity-based method. (b) 2D depiction of `‘HME’` with a Depiction Penalty Score of 0, indicating a high-quality representation with no bond clashes generated using the template-based method.

### Identification of scaffolds and fragments

Medicinal chemists widely use the concept of classifying compounds based on their molecular scaffolds to group molecules with similar properties [45]. Figure 5a showcases the scaffold identified by PDBe CCDUtils using the BRICS fragmentation rule [39] for the CCD component CVV when bound to the human kappa opioid receptor (PDB entry 6b73).

**Figure 5.**
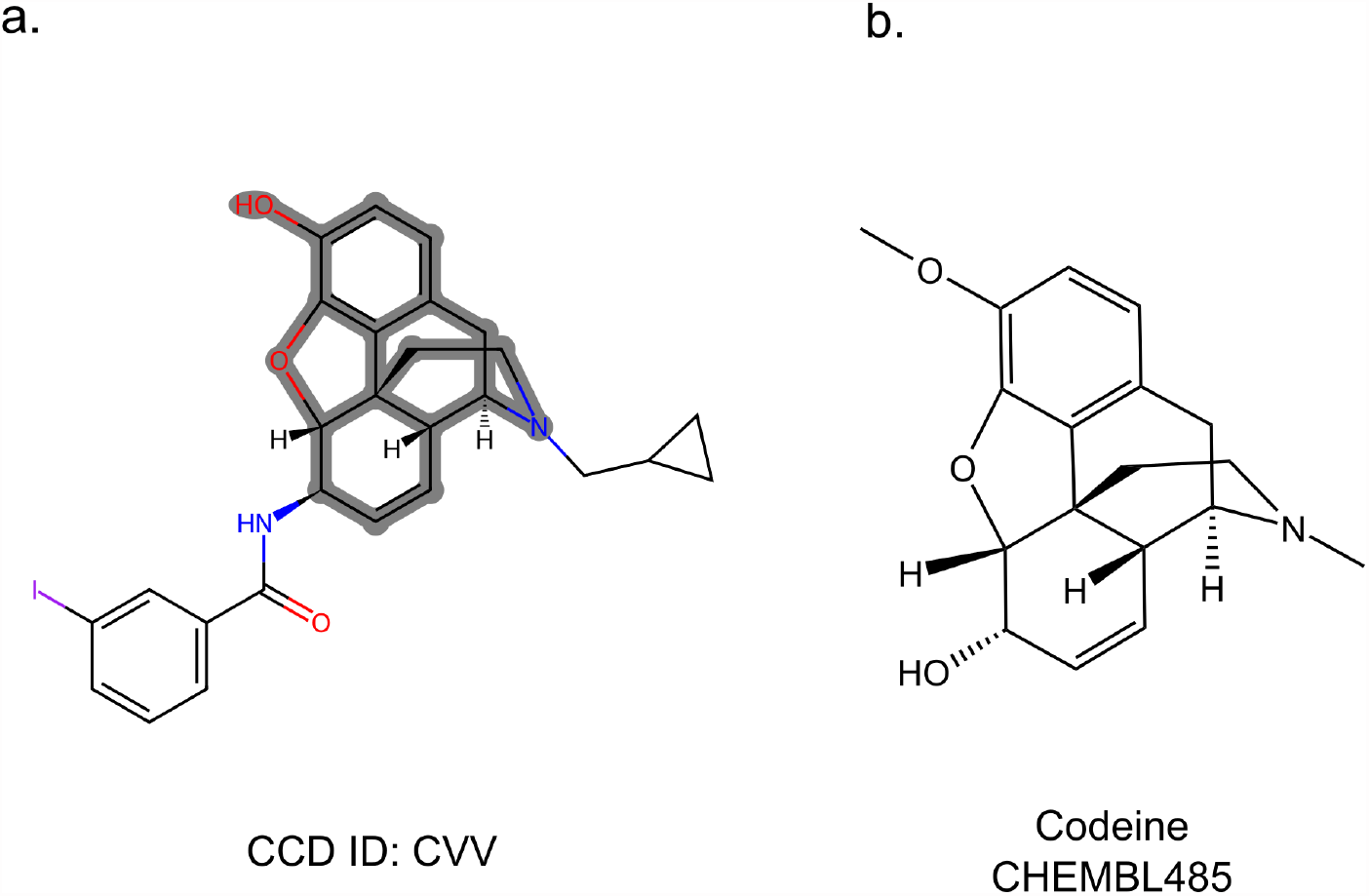
Example for identification of scaffolds. (a) 2D depiction of CCD component CVV highlighted with scaffold identified by PDBe CCDUtils. (b) 2D depiction of Codeine.

Interestingly, it is an exact match to the scaffold of Codeine (ChEMBL485), depicted in Figure 5b. Codeine is a known analgesic that targets various opioid receptors [46], and its biological activity is well-documented in ChEMBL [47]. Although the PDB does not contain the structure of Codeine, the shared scaffold between Codeine and CCD component CVV suggests that Codeine may interact with the Human kappa opioid receptor in a similar manner to CVV.

Furthermore, the PDBe CCDUtils’ fragment library was used to identify matching fragments to CVV, and these are listed in Table 2. This process allows researchers to explore smaller molecular fragments that can potentially interact with the target of interest and serve as starting points for drug development [48]. By understanding the shared scaffolds and fragments, medicinal chemists can make informed decisions and design new compounds with improved pharmacological properties and higher chances of success in drug discovery[49]

**Table 2.**
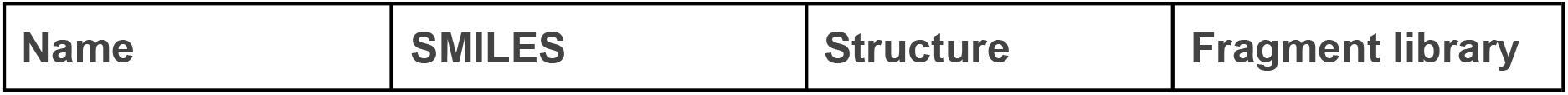
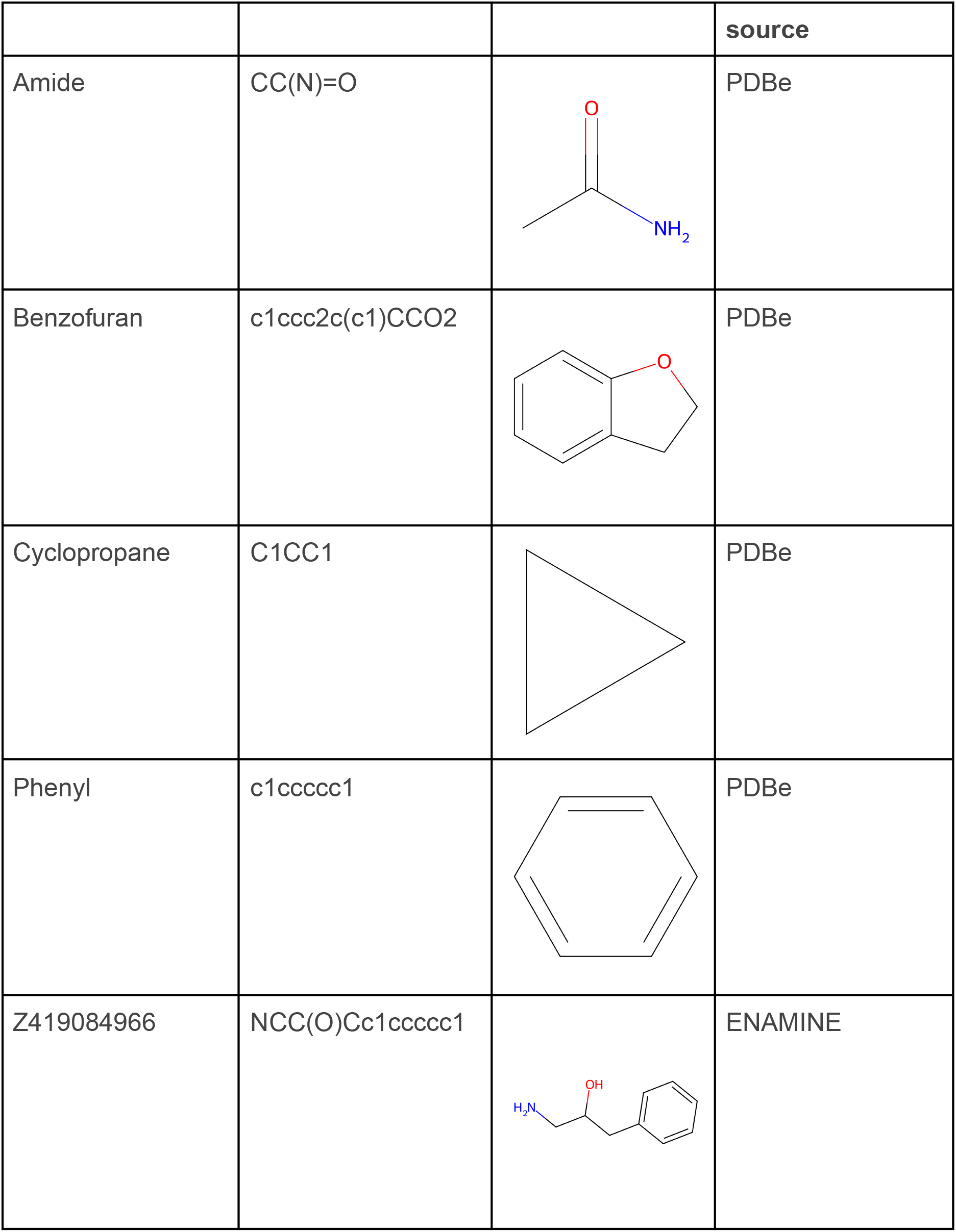
Matching fragments of CVV from PDBe CCDUtils’ fragment library.

| Name | SMILES | Structure | Fragment library |
| --- | --- | --- | --- |

**Table 2. Matching fragments of CVV from PDBe CCDUtils' fragment library**
|  |  |  | source |
| --- | --- | --- | --- |
| Amide | CC(N)=O |  | PDBe |
| Benzofuran | c1ccc2c(c1)CCO2 |  | PDBe |
| Cyclopropane | C1CC1 |  | PDBe |
| Phenyl | c1ccccc1 |  | PDBe |
| Z419084966 | NCC(O)Cc1ccccc1 |  | ENAMINE |

### Additional small molecule related meta-data

PDBe CCDUtils serves as the primary package in the PDBeChem pipeline, a weekly update process for the small molecule information in PDB [26]. The pipeline processes three key PDBx/mmCIF reference files related to small molecules: CCDs, PRDs and CLCs, enriching them with valuable additional information. This additional information encompasses RDKit-generated 3D conformers, 2D coordinates of the ligands, RDKit-generated physicochemical properties and details about scaffolds and fragments. Moreover, the pipeline also establishes cross-references to other external small molecule databases like ChEMBL, PubChem, KEGG and DrugBank through UniChem mapping. Additional information on descriptions, synonyms, taxonomy, and known targets are added for the small molecules mapped to DrugBank. Synonyms collected from ChEMBL and wwwPDB are also added to the updated PDBx/mmCIF files. All the files generated by the PDBeChem pipeline are available from the PDBe FTP area [24]. This directory contains a detailed readme file elucidating the contents, along with separate folders for each enriched small molecule reference file: CCD, PRD, and CLC. Each CCD/PRD or CLC identifier has this data in various types of files, further elaborated in Table 3.

**Table 3.**
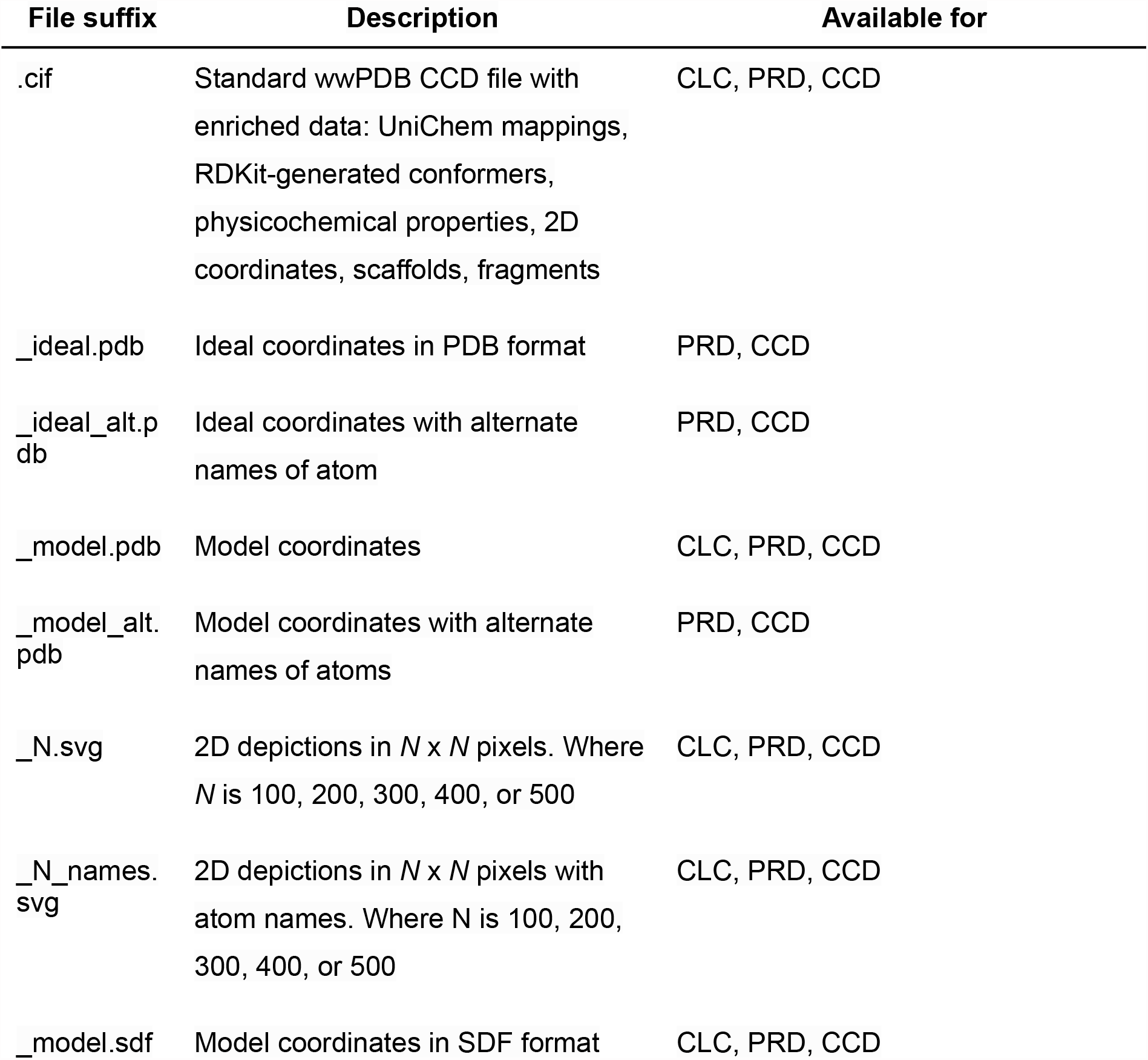
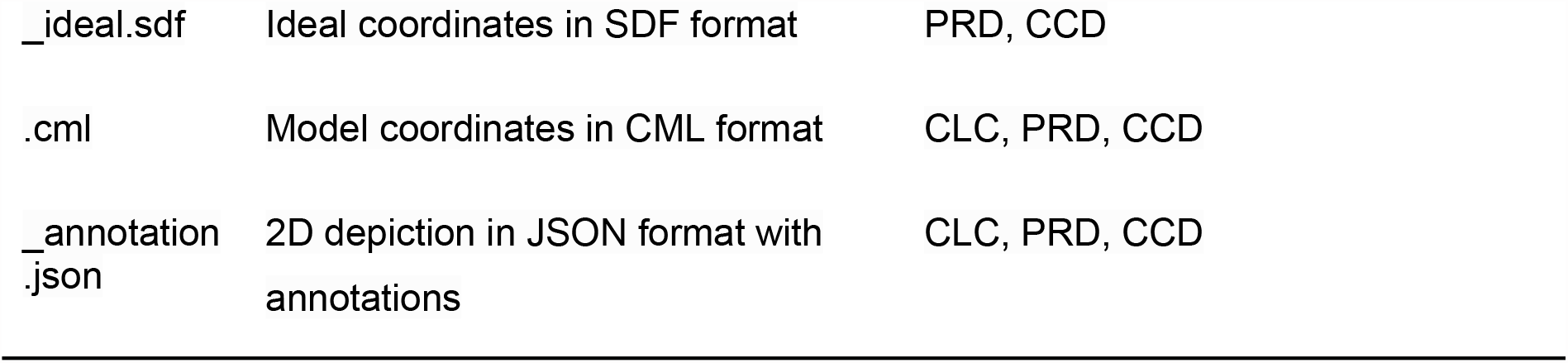
Details of enrichments and associated files for CCD, PRD or CLC identifiers generated by the PDBeChem pipeline using PDBe CCDUtils.

## Conclusions

PDBe CCDUtils is a versatile Python package designed to streamline the analysis and manipulation of small molecules in the Protein Data Bank (PDB) by expanding the core functionalities of RDKit. This tool addresses the challenges faced by researchers working with a diverse range of small molecules in the PDB, offering an accessible and efficient toolkit tailored to meet the needs of researchers in cheminformatics, structural biology, bioinformatics, and computational chemistry.

The integration of RDKit with PDBe CCDUtils allows users to leverage the powerful features provided by RDKit while offering additional enhancements, such as the definition of CLCs for large, complex ligands, which are often split into individual CCDs in the PDB. This approach enables a more accurate representation and analysis of complex ligands in biological systems, with the added functionality of generating unique InChIKeys for each CLC molecule.

## Availability and requirements

**Project name:** ccdutils

**Project home page:** https://github.com/PDBeurope/ccdutils

**Operating systems:** Platform independent

**Programming language:** Python

**Other requirements:** Python 3.9+; RDKit `2022.09.4` or higher

**Licence:** Apache 2.0

**Any restrictions to use by non-academics:** see the licence

**Tutorials:** https://pdbeurope.github.io/ccdutils/guide/intro.html

## Authors’ contributions

I.R.K. designed and implemented the code and worked on the original draft. P.C. managed the project and worked on the original draft. L.P., O.S.S., N.N., Q.Y. designed and implemented the PDBe CCDUtils code. S.A. and S.S. worked on setting up CI/CD for CCDUtils and running it as part of the PDBe weekly release process. M.V. coordinated the project, worked on the original draft and contributed to securing funding. S.V. contributed to conceptualization, supervision, and secured funding. All authors contributed to the manuscript.

## References

1. wwPDB consortium, Burley SK, Berman HM, et al (2019) Protein Data Bank: the single global archive for 3D macromolecular structure data. Nucleic Acids Res 47:D520–D528. https://doi.org/10.1093/nar/gky949

2. Berman H, Henrick K, Nakamura H (2003) Announcing the worldwide Protein Data Bank. Nat Struct Biol 10:980. https://doi.org/10.1038/nsb1203-980

3. Peat TS, Christopher JA, Newman J (2005) Tapping the Protein Data Bank for crystallization information. Acta Crystallogr D Biol Crystallogr 61:1662–1669. https://doi.org/10.1107/S0907444905033202

4. Caffrey M, Cherezov V (2009) Crystallizing membrane proteins using lipidic mesophases. Nat Protoc 4:706–731. https://doi.org/10.1038/nprot.2009.31

5. McPherson A, Cudney B (2014) Optimization of crystallization conditions for biological macromolecules. Acta Crystallogr Sect F Struct Biol Commun 70:1445–1467. https://doi.org/10.1107/S2053230X14019670

6. Garman E (2003) “Cool” crystals: macromolecular cryocrystallography and radiation damage. Curr Opin Struct Biol 13:545–551. https://doi.org/10.1016/j.sbi.2003.09.013

7. Pflugrath JW (2015) Practical macromolecular cryocrystallography. Acta Crystallogr Sect F Struct Biol Commun 71:622–642. https://doi.org/10.1107/S2053230X15008304

8. Jang K, Kim HG, Hlaing SHS, et al (2022) A Short Review on Cryoprotectants for 3D Protein Structure Analysis. Crystals 12:. https://doi.org/10.3390/cryst12020138

9. Mukhopadhyay A, Borkakoti N, Pravda L, et al (2019) Finding enzyme cofactors in Protein Data Bank. Bioinformatics 35:3510–3511. https://doi.org/10.1093/bioinformatics/btz115

10. Westbrook J, Burley SK (2019) How Structural Biologists and the Protein Data Bank Contributed to Recent US FDA New Drug Approvals. Struct Lond Engl 1993 27:211–217. https://doi.org/10.1016/j.str.2018.11.007

11. Vetting MW, Al-Obaidi N, Zhao S, et al (2015) Experimental Strategies for Functional Annotation and Metabolism Discovery: Targeted Screening of Solute Binding Proteins and Unbiased Panning of Metabolomes. Biochemistry 54:909–931. https://doi.org/10.1021/bi501388y

12. Goodsell DS, Zardecki C, Di Costanzo L, et al (2020) RCSB Protein Data Bank: Enabling biomedical research and drug discovery. Protein Sci Publ Protein Soc 29:52–65. https://doi.org/10.1002/pro.3730

13. Westbrook JD, Shao C, Feng Z, et al (2015) The chemical component dictionary: complete descriptions of constituent molecules in experimentally determined 3D macromolecules in the Protein Data Bank. Bioinformatics 31:1274–1278. https://doi.org/10.1093/bioinformatics/btu789

14. Sen S, Young J, Berrisford JM, et al (2014) Small molecule annotation for the Protein Data Bank. Database J Biol Databases Curation x2014:bau116. https://doi.org/10.1093/database/bau116

15. wwPDB Deposition Policies and wwPDB Biocuration Procedures. http://www.wwpdb.org/documentation/procedure#toc_4

16. Dutta S, Dimitropoulos D, Feng Z, et al (2014) Improving the representation of peptide-like inhibitor and antibiotic molecules in the Protein Data Bank. Biopolymers 101:659–668. https://doi.org/10.1002/bip.22434

17. Callaway E (2015) The revolution will not be crystallized: a new method sweeps through structural biology. Nature 525:172–174. https://doi.org/10.1038/525172a

18. Behzadi P, Gajdács M (2022) Worldwide Protein Data Bank (wwPDB): A virtual treasure for research in biotechnology. Eur J Microbiol Immunol 11:77–86. https://doi.org/10.1556/1886.2021.00020

19. Future Planning: PDB entries with extended CCD or PDB IDs will be distributed in the PDBx/mmCIF format only. https://www.wwpdb.org/news/news?year=2022#630fee4cebdf34532a949c34

20. RDKit: Open-source cheminformatics. https://www.rdkit.org/. Accessed 6 Jun 2023

21. Westbrook JD, Young JY, Shao C, et al (2022) PDBx/mmCIF Ecosystem: Foundational Semantic Tools for Structural Biology. J Mol Biol 434:167599. https://doi.org/10.1016/j.jmb.2022.167599

22. The wwPDB CCD in mmCIF format. https://files.wwpdb.org/pub/pdb/data/monomers/components.cif

23. The wwPDB PRD in mmCIF format. https://ftp.wwpdb.org/pub/pdb/data/bird/prd/

24. PDBeChem FTP Area-Enhanced small molecule data in PDB. http://ftp.ebi.ac.uk/pub/databases/msd/pdbechem_v2

25. Wojdyr M (2022) GEMMI: A library for structural biology. J Open Source Softw 7:4200. https://doi.org/10.21105/joss.04200

26. Armstrong DR, Berrisford JM, Conroy MJ, et al (2020) PDBe: improved findability of macromolecular structure data in the PDB. Nucleic Acids Res 48:D335–D343. https://doi.org/10.1093/nar/gkz990

27. 3D Structure Generator CORINA Classic. https://mn-am.com/. Accessed 7 Jun 2023

28. OMEGA 4.2.2.0. OpenEye, Cadence Molecular Sciences, santa Fe, NM. https://www.eyesopen.com. Accessed 7 Jun 2023

29. Hagberg AA, Schult DA, Swart PJ (2008) Exploring Network Structure, Dynamics, and Function using NetworkX. In: Varoquaux G, Vaught T, Millman J (eds) Proceedings of the 7th Python in Science Conference. Pasadena, CA USA, pp 11–15

30. Landrum G (2023) Drawing options explained. In: RDKit Blog. https://greglandrum.github.io/rdkit-blog/posts/2023-05-26-drawing-options-explained.html. Accessed 7 Jun 2023

31. Kim S, Chen J, Cheng T, et al (2023) PubChem 2023 update. Nucleic Acids Res 51:D1373–D1380. https://doi.org/10.1093/nar/gkac956

32. The PDBe CCDUtils hand-curated templates for 2D image generation of ligands in PDB. https://github.com/PDBeurope/ccdutils/tree/master/pdbeccdutils/data/general_templates

33. Kim S, Thiessen PA, Cheng T, et al (2018) An update on PUG-REST: RESTful interface for programmatic access to PubChem. Nucleic Acids Res 46:W563–W570. https://doi.org/10.1093/nar/gky294

34. Strang G (2016) Introduction to Linear Algebra. Wellesley

35. Virtanen P, Gommers R, Oliphant TE, et al (2020) SciPy 1.0: fundamental algorithms for scientific computing in Python. Nat Methods 17:261–272. https://doi.org/10.1038/s41592-019-0686-2

36. Manelfi C, Gemei M, Talarico C, et al (2021) “Molecular Anatomy”: a new multi-dimensional hierarchical scaffold analysis tool. J Cheminformatics 13:54. https://doi.org/10.1186/s13321-021-00526-y

37. Kim J, Kim H, Park SB (2014) Privileged Structures: Efficient Chemical “Navigators” toward Unexplored Biologically Relevant Chemical Spaces. J Am Chem Soc 136:14629–14638. https://doi.org/10.1021/ja508343a

38. Bemis GW, Murcko MA (1996) The Properties of Known Drugs. 1. Molecular Frameworks. J Med Chem 39:2887–2893. https://doi.org/10.1021/jm9602928

39. Degen J, Wegscheid-Gerlach C, Zaliani A, Rarey M (2008) On the Art of Compiling and Using “Drug-Like” Chemical Fragment Spaces. ChemMedChem 3:1503–1507. https://doi.org/10.1002/cmdc.200800178

40. Kirsch P, Hartman AM, Hirsch AKH, Empting M (2019) Concepts and Core Principles of Fragment-Based Drug Design. Molecules 24:4309. https://doi.org/10.3390/molecules24234309

41. Grygorenko OO (2021) Enamine Ltd.: The Science and Business of Organic Chemistry and Beyond. Eur J Org Chem 2021:6474–6477. https://doi.org/10.1002/ejoc.202101210

42. Cox OB, Krojer T, Collins P, et al (2016) A poised fragment library enables rapid synthetic expansion yielding the first reported inhibitors of PHIP(2), an atypical bromodomain. Chem Sci 7:2322–2330. https://doi.org/10.1039/C5SC03115J

43. The PDBeCCDUtils fragment library manually curated by PDBe, ENAMINE and Diamond-SGC-iNext Poised Library (DSiP). https://github.com/PDBeurope/ccdutils/tree/master/pdbeccdutils/data/general_template s

44. Gally J-M, Pahl A, Czodrowski P, Waldmann H (2021) Pseudonatural Products Occur Frequently in Biologically Relevant Compounds. J Chem Inf Model 61:5458–5468. https://doi.org/10.1021/acs.jcim.1c01084

45. Medina-Franco JL, Flores-Padilla EA, Chávez-Hernández AL (2022) Chapter 23 - Discovery and development of lead compounds from natural sources using computational approaches. In: Mukherjee PK (ed) Evidence-Based Validation of Herbal Medicine (Second Edition). Elsevier, pp 539–560

46. McDonald J, Lambert D (2015) Opioid receptors. BJA Educ 15:219–224. https://doi.org/10.1093/bjaceaccp/mku041

47. Mendez D, Gaulton A, Bento AP, et al (2019) ChEMBL: towards direct deposition of bioassay data. Nucleic Acids Res 47:D930–D940. https://doi.org/10.1093/nar/gky1075

48. Price AJ, Howard S, Cons BD (2017) Fragment-based drug discovery and its application to challenging drug targets. Essays Biochem 61:475–484. https://doi.org/10.1042/EBC20170029

49. Sadybekov AV, Katritch V (2023) Computational approaches streamlining drug discovery. Nature 616:673–685. https://doi.org/10.1038/s41586-023-05905-z

